# Generation of Mouse-Zebrafish Hematopoietic Tissue Chimeric Embryos for Hematopoiesis and Host-Pathogen Interaction Studies

**DOI:** 10.1101/216895

**Authors:** Margarita Parada-Kusz, Anne Clatworthy, Elliott J. Hagedorn, Cristina Penaranda, Anil V. Nair, Jonathan E. Henninger, Christoph Ernst, Brian Li, Raquel Riquelme, Humberto Jijon, Eduardo J. Villablanca, Leonard I. Zon, Deborah Hung, Miguel L. Allende

## Abstract

Xenografts of the hematopoietic system are extremely useful as disease models and for translational research. Zebrafish xenografts have been widely used to monitor blood cancer cell dissemination and homing due to the optical clarity of embryos and larvae, which allow unrestricted *in vivo* visualization of migratory events. To broaden the scope of xenotransplantation studies in zebrafish, we have developed a technique that transiently generates hematopoietic tissue chimeras by transplanting murine bone marrow cells into zebrafish blastulae. This procedure leads to mammalian cell integration into the fish developmental hematopoietic program. Monitoring zebrafish chimeras at different time points post fertilization using *in vivo* time-lapse and confocal imaging showed murine cell co-localization with developing primitive and definitive hematopoietic tissues, intravasation into fish circulation, and dynamic hematopoietic cell-vascular endothelial and hematopoietic cell-niche interactions. Immunohistochemistry assays performed in chimeric animals showed that, after engraftment, murine cells expressed antigens related to i) hematopoietic stem and progenitor cells, ii) active cell proliferation, and iii) myeloid cell lineages. Lastly, xenografted zebrafish larvae infected with *Klebsiella pneumoniae* showed murine immune cells trafficking to bacterial foci and interacting with bacterial cells. Overall, these results show that mammalian bone marrow cells xenografted in zebrafish integrate into the host hematopoietic system revealing highly conserved molecular mechanisms of hematopoiesis between zebrafish and mammals. In addition, this procedure introduces a useful and simple method that improves and broadens the scope of hematopoietic tissue xenotransplantation studies in zebrafish.

## BACKGROUND

Xenotransplantation has been a powerful tool for understanding mammalian cancer, hematopoiesis, immunity and infectious diseases. For example, humanized mice, where animals are engrafted with human cord blood-derived hematopoietic stem cells or human adult peripheral blood mononuclear cells, allow for analysis of the cellular and molecular processes that mediate human hematopoietic and immune cell interactions within stromal niches and with pathogens *in vivo*^1–4^. Furthermore, xenotransplants offer the unique opportunity to study the function of human disease-associated single nucleotide polymorphisms that are non-existent or irreproducible in other species. Current research, however, is limited by the challenges of quantitatively measuring and tracking individual cell responses to these complex events^5–7^. Observing cellular interactions in real time would allow the identification and precise evaluation of key processes between various cells and tissues that promote or restrict responses at the appropriate time and location. Intravital microscopy has been developed to perform these analyses in mouse models but lacks resolution and often requires more invasive follow-up procedures that can interfere with normal cell behaviors. Zebrafish embryos and larvae, in contrast, are transparent, making them ideally suited to perform *in vivo* analyses in unperturbed live animals.

Strong conservation of genes and biological processes between zebrafish and mammals has made zebrafish a well-established model for basic research of the hematopoietic and innate immune systems^8–16^. Xenotransplantation assays have allowed the model to be used as an inexpensive platform for assessing cancer cell behavior and to perform drug screens *in vivo* with translational applications^17–24^. Recently, xenotransplantation of human CD34+ cells and multiple myeloma cells into the blood stream of zebrafish embryos evidenced that human cells disseminate to the caudal hematopoietic tissue and actively respond to the hematopoietic niche^25–26^. In a similar context, xenotransplantation of human macrophages showed these cells can survive and acquire an activated phenotype in the zebrafish^27^. These results demonstrate the scientific and clinical potential of zebrafish blood cell xenotransplantation studies. For example, mammalian-zebrafish blood chimeric embryos could be used for visualization of hematopoietic cell-niche interactions and immune cell participation in bacterial infection. Furthermore, since some pathogens are species-specific and mammalian cells are unique, among other things, in the repertoire of antimicrobial peptides that they express upon infection^28–29^, mammalian-zebrafish blood chimeras could expand the utility of zebrafish as an infection model. However, the ability of xenografted mammalian blood cells to respond to bacterial pathogens or other stimuli in fish, to the best of our knowledge, has not been addressed yet. In addition, the scope of these experiments is limited by the cumbersome transplantation procedures that limit the number of cells that can be transplanted and the number of chimeric animals that can be generated.

Here, we improve and extend the range of zebrafish blood cell xenotransplantation analyses by developing a fast and efficient method that generates a high number of transient chimeric zebrafish embryos with mammalian hematopoietic cells. This technique is based upon injection of murine bone marrow cells into zebrafish blastulae, which leads to mammalian cell integration into the fish hematopoietic developmental program. As proof of concept, we illustrate the value of mouse-zebrafish chimeras by showing non-invasive real time visualization of murine cell migration and homing to primitive and definitive hematopoietic tissues, intravasation into fish circulation, hematopoietic cell-vascular endothelial and hematopoietic cell-niche interactions, as well as xenotransplanted immune cell response to bacterial infection. This straightforward methodology can be scaled up to allow rapid and efficient assays for the evaluation of genetic or pharmacological interventions on mammalian cells and for discovery of novel processes related to mammalian hematopoiesis and immune cell dynamics.

## MATERIALS AND METHODS

### Animal use and generation of mouse-zebrafish chimeras

Wild-type AB, *nacre*^30^ mutants and *ubi:mcherry*^31^, *flk:dsred*^32^*, mpeg1:mcherry*^33^ and *mpx:gfp*^34^ transgenic embryos were raised and staged according to Westerfield^35^. Zebrafish studies were approved by the Massachusetts General Hospital Institutional Animal Care and Use Committee.

### Isolation of mouse bone marrow stem and progenitor cells and neutrophils

Mouse bone marrows cells were isolated from C57BL/6 or CByJ.B6-Tg(UBC-GFP)30Scha/J (UBI-GFP) animals. For enrichment of stem and progenitor cells or neutrophils, bone marrow cell suspensions were incubated with lineage cell depletion kit or neutrophil isolation kit from Miltenyi Biotec (Auburn, CA). The procedure was conducted according to the manufacturer’s recommendations, with one modification involving addition of an extra 10 μL of anti-TER-119 (Ebioscience, #13-5921-81) to the antibody cocktail.

### Human cell lines

Urinary bladder epithelial T24 (ATCC^®^ HTB-4™) cells, the neuroblastoma cell line SH-SY5Y (ATCC^®^ CRL-2266™) and promyelocytic HL-60 (ATCC^®^ CCL-240™) cells were cultured according to the manufacturer’s recommendations.

### Transplantation procedure

Isolated mouse cells were microinjected directly into the blastoderm of 3–5 hpf zebrafish blastulae with a range of 500 to 5,000 cells. Transplanted cells were quantified by injecting into PBS and subsequent analysis by flow cytometry. For a complete transplantation protocol description see **Supplemental Note S1**.

### Cell labeling

For labeling and *in vivo* visualization, cells were incubated with the vital dye CellTrace^TM^ Violet Cell Proliferation Kit (450 nm blue emission) (Thermo Fisher Scientific Inc., #C34557) or CellTrace^TM^ CSFE (517 nm green emission) (Thermo Fisher Scientific Inc., #C34554) according to the manufacturer’s recommendations.

### Infection and bacterial strains

A clinical isolate of *Klebsiella pneumoniae* was used for infection experiments. For *in vivo* visualization, the bacteria were transformed with the pRSET-tdTomato vector, where tdTomato is constitutively expressed^36^. Bacteria were cultured over night at 37°C in LB media with or without carbenicillin (for transformed cells selection). For infections, embryos were anesthetized with tricaine^35^, immobilized in 1% low–melting point agarose (Sigma) and covered with E3 medium, both containing tricaine. Larvae were microinjected into the tail muscles or the otic vesicle. Bacterial inocula were determined by injecting into PBS and later plating into LB agar plates. After infection, the xenotransplanted animals were incubated at 30°C to favor the mammalian cell response.

### Whole-mount immunohistochemistry

Wild type and chimeric zebrafish larvae of 2–3 dpf were euthanized with tricaine^35^ in E3 medium, fixed in fresh 4% paraformaldehyde in PBST (PBS, 0.3% Triton X-100) and dehydrated using methanol (1 hour at −20°C). Embryos were rehydrated using decreasing concentrations of methanol in PBST, and permeabilized using 1 µg/mL proteinase K at room temperature for 1 hour. Larvae were then re-fixed and incubated with a cold ethanol/acetone solution (2:1) for 10 minutes at −20°C. Fixed and permeabilized embryos were then incubated in blocking solution (10 mg/mL BSA, 2% BFS, 1% DMSO, 0.1% Triton X-100) and later incubated either with the anti-c-kit against mouse/rat biotin conjugated antibody (Abcam, #ab25022), the anti-mouse Ly-6G/Ly-6C (Gr-1) biotin conjugated antibody (Biolegend, #108404), the anti-mouse Ki-67-FITC (Ebioscience, #11-5698-80) or the anti-mouse F4/80-FITC antibody (Ebioscience, #11-4801-82). Streptavidin-FITC (Ebioscience, #11-4317-87) was used as secondary antibody for biotin conjugated primary antibodies. For quantification, the CHT of chimeric animals was visualized and manually counted for positive murine cells for a specific antibody on an epifluorescence microscope (Zeiss Axio Observer). Percentage was calculated with respect to the total murine CHT cell count in individual larvae (blue labeled cells). Mean and standard error was calculated from total larvae analyzed.

### TUNEL assays

Wild type and chimeric zebrafish larvae of 2–3 dpf were euthanized, fixed, permeabilized, and refixed as described for whole mount immunohistochemistry. For the Terminal deoxynucleotidyl transferase (TdT) dUTP Nick-End Labeling (TUNEL) assay, the ApopTag^®^ Red In Situ Apoptosis Detection Kit (Milipore Sigma, #S7165) was used. Briefly, fixed and permeabilized embryos were incubated in equilibration buffer for 1 hour at room temperature, and then incubated in a reaction mix (20 μL of equilibration buffer, 12 μL of reaction buffer, 6 μL of TdT enzyme, 0.5 μL of 10% Triton X-100) overnight at 37°C. Larvae were then washed several times with PBST and then incubated in blocking solution (10 mg/mL BSA, 2% BFS, 1% DMSO, 0.1% Triton X-100). Anti-Digoxigenin-Rhodamine (Roche, #11207750910) was used as conjugated primary antibody. Quantification was performed as described above.

### Time-lapse confocal fluorescence imaging of live zebrafish embryos and larvae

Embryos were anesthetized with tricaine^35^, immobilized in 1% low–melting point agarose (Sigma) containing tricaine on 35-mm glass-bottom dishes (MatTek), covered with E3 medium containing tricaine and imaged on an epifluorescence microscope (Zeiss Axio Observer), an A1R or C2 (Nikon) confocal microscope, or an Eclipse Ti (Nikon) spinning disk confocal microscope as previously described^37^. All images were adjusted for brightness and contrast to improve visualization.

### Flow cytometry analyses

Murine pre- and post-enrichment bone marrow cells were incubated with LIVE/DEAD™ Fixable Near-IR Dead Cell Stain (Thermo Fisher Scientific Inc., #L34975) and the anti-CD45-FITC (Biolegend, #103107), anti-mouse c-kit-APC (Biolegend, #105811) and anti-mouse CD11b-PerCP/Cy5.5 (Biolegend, #101227) primary antibodies. Samples were then analyzed in a NovoCyte flow cytometer (ACEA Biosciences Inc.). For zebrafish chimeras-cell analyses, embryos were euthanized with tricaine^35^ in E3 medium (8-12 embryos per condition) and mechanically dissociated by sterile razor blades. A single cell suspension was prepared by collecting dissociated tissue in 0.5 mL 0.9X PBS/2% FBS and passing the sample through a 40-µM nylon mesh. Samples were subjected to 3 nM DRAQ-7 Dead Cell Stain (Abcam, #ab109202) and analyzed by flow cytometry on a BD FACSAria II (Becton Dickinson).

## RESULTS

### Generation of mouse-zebrafish hematopoietic tissue chimeric embryos

The method developed here is based upon: (1) isolation of mouse bone marrow cells, (2) enrichment for hematopoietic stem and progenitor cells (HSPCs), (3) fluorescent labeling, and (4) transplantation into the blastoderm of zebrafish embryos (See methods and Supplementary Note). First, bone marrow cells are isolated from both femurs and tibiæ from 1 mouse. Bone tissue is homogenized and marrow cells are collected and incubated with an antibody cocktail in order to enrich for lineage-negative cells (HSPCs) by means of negative selection. Analysis of the cell population obtained determined that around 50% of cells are c-kit+ and ~39% are double c-kit+/CD11b+ (Supplementary Figure S1). After enrichment, cells are stained for *in vivo* monitoring with a blue fluorescent emission vital dye and later injected directly into the blastoderm of zebrafish embryos at 3–5 hours post fertilization (hpf) (Figure 1A). To facilitate murine cell tracking and visualization at subsequent stages, *nacre*^−/−^ zebrafish mutants with impaired pigmentation development^30^ were used as hosts. In our hands, up to 800 embryos can be transplanted in 3 hours by one person, leading to around 500 viable embryos the following day. Strikingly, embryos can be injected with a range of approximately 500 to up to 5,000 cells with less than 5% of viable chimeric animals showing morphological abnormalities at 3 days post fertilization (dpf) (Figure 1B).

**Figure 1.**
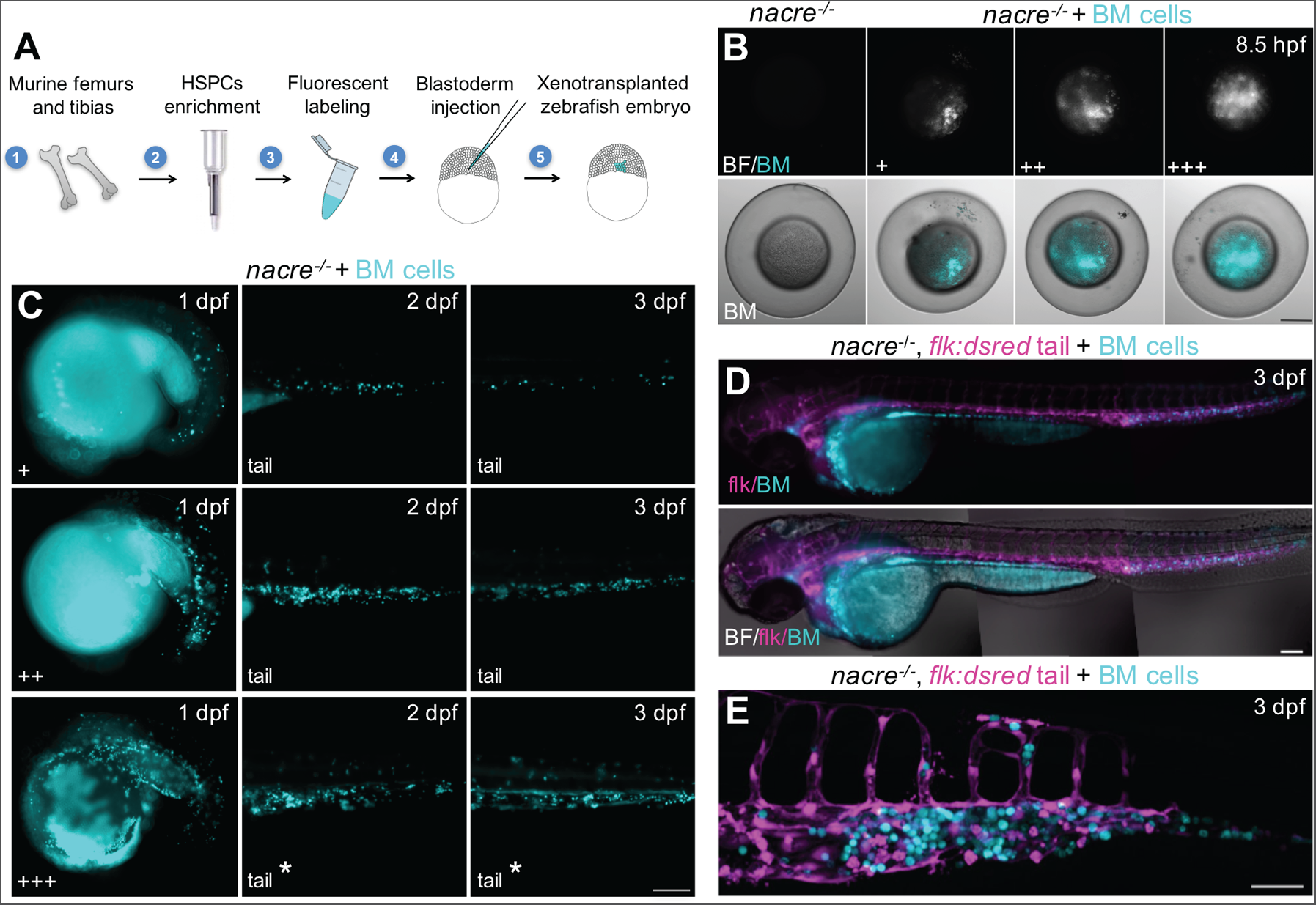
Generation of hematopoietic tissue chimeras by transplantation of mouse bone marrow cells into zebrafish blastulae. A) Experimental procedure. Bone marrow cells are isolated, enriched for hematopoietic stem and progenitor cells (HSPCs) by means of negative selection, blue fluorescently labeled, and transplanted into the blastoderm of 3–5 hpf zebrafish embryos. B) Representative epifluorescence images (animal pole views) of zebrafish embryos transplanted with 3 different quantities of injected mouse bone marrow cells (+, ++ and +++, respectively, within a range of ~500 to 5,000 cells). Scale bar: 100 µm. C) Representative images of chimeric embryos showing mouse cells in the ICM at 1 dpf (left column), in the PBI and AGM at 2 dpf (middle column), and in the CHT at 3 dpf (right column). Quantification data was obtained from 2 independent experiments including more than 500 animals. Asterisks indicate animals with cells into circulation. Scale bar: 200 µm. D) Pseudo-colored epifluorescence images of a global view of a xenotransplanted *flk:dsred* fish at 3 dpf (mouse cells in blue; fish vasculature in magenta). Individual images from fish head, trunk and tail were joined to create a whole embryo high magnification image. Scale bar: 100 µm. E) Pseudo-colored confocal image of the tail region of a xenotransplanted *flk:dsred* fish at 3 dpf. Scale bar: 50 µm. BF: bright field. BM cells: mouse bone marrow cells.

We examined chimeric animals throughout development. Visualization at 1 dpf evidenced transplanted cells distributing over the yolk sac and the whole body of the embryo, with 67.8% of chimeric animals showing cells within the fish intermediate cell mass (ICM) (Figure 1C, left column). By transplanting murine cells into the transgenic endothelial vasculature reporter line *flk:dsred*^32^, we determined that 83.2% of chimeric animals had murine cells residing within the fish posterior blood island (PBI) and aorta-gonad-mesonephros (AGM) at 2 dpf (Figure 1C, middle column). At 3 dpf, 93.3% had murine cells in the caudal hematopoietic tissue (CHT, Figure 1C, right column, and 1D). We also observed a small fraction of chimeric animals with murine cells neighboring the developing thymic lobes at 3 dpf, suggesting that murine cells can colonize the developing thymus, a definitive hematopoietic organ in zebrafish^38^ (Supplemental Figure S2). Taken together, these results suggest active homing of murine bone marrow cells into the developing fish hematopoietic niche.

To evaluate if trafficking to the CHT could be visualized for other mammalian cell types transplanted into the blastoderm of zebrafish, or if it is rather a unique process for cells of the hematopoietic lineage, we xenotransplanted murine neutrophils, human neuroblastoma cells, human bladder epithelial cells and human promyelocytic cells. Visualization of chimeric animals show neuroblastoma cells localizing mainly within the head region, epithelial cells localizing around the ventral anterior part of the fish body, and neutrophils and promyelocytic cells colonizing the CHT (Supplemental Figures S3–S5). These results suggest that engrafting of the zebrafish CHT is a specific process for cells of hematopoietic lineage, and indicate that human cells can also be engrafted and followed with this procedure.

We next evaluated whether we could distinguish different murine cell types residing in the zebrafish hematopoietic tissue. Confocal imaging showed that murine bone marrow cells have similar patterns of intracellular granules and morphology to fish cells in the CHT (Figure 1E and Supplemental Figure S6). Immunohistochemistry assays performed in 2 dpf chimeric embryos showed that 35.6% of murine cells express the c-kit protein, which is highly expressed in hematopoietic stem and progenitor cells, and that 21.1% of cells express the ki-67 protein related to active cell proliferation (Supplemental Figures S7 and S8). A few chimeric embryos displayed morphological abnormalities. Among these, we observed animals with an enlarged CHT containing varying levels of blue fluorescence emission (Supplemental Figure S9). Since the vital dye utilized to track murine cells in our experiments becomes diluted between mother and daughter cells upon cell division, the observed heterogeneous levels of fluorescence suggest that unregulated or excessive local murine cell proliferation can lead to deformation of the host niche. In addition, evaluation of myeloid cell lineage markers revealed that 32.5% of murine cells in the hematopoietic niche expressed the granulocyte cell associated Gr1 protein, and 13.4% of cells expressed the monocyte-macrophage F4/80 protein (Supplemental Figures S10 and S11). Altogether, these results indicate that the CHT of chimeric animals contains proliferating murine hematopoietic progenitors as well as myeloid lineage cells.

### Xenotransplantation of murine cells into zebrafish blastulae does not induce vital dye transfer or cell fusion events

To exclude the possibility that the vital dye used to stain the murine cells could be transferring to fish cells, or that murine cells may be fusing with fish cells after xenotransplantation, we transplanted blue-stained bone marrow cells from UBI-GFP transgenic (ubiquitously green) mice into *ubi:mcherry*^31^ transgenic fish. These chimeras have green and blue-labeled murine cells in a ubiquitously red fluorescent zebrafish host. Confocal imaging of an 8 hpf and a 20 hpf transplanted animal showed no triple-color labeled cells (white emission), suggesting negligible dye transfer or cell fusion events (Figure 2A). To corroborate these results, cells from 2 dpf chimeric animals were analyzed by flow cytometry. Examination of the physical parameters of live pre-gated cells showed murine bone marrow cells as a uniform population that are, on average, slightly larger than fish cells (Figure 2B). Cell-color emission analysis detected scarce cells showing double green and red label (0.017%), as well as few cells double labeled blue and red (0.45%) (Figure 2C–D). In addition, analysis of green and blue-labeled cells demonstrated that all green cells are blue, indicating that all transplanted cells retain the blue stain during this timeframe (Figure 2E). Finally, to test for the existence of triple colored cells, all green and blue-labeled cells were tested for red emission, showing that only a small fraction (0.87%) are triple labeled (Figure 2F). These results indicate that transplantation of stained murine cells does not lead to significant vital dye transfer to fish cells and that cell fusion events are rare.

**Figure 2.**
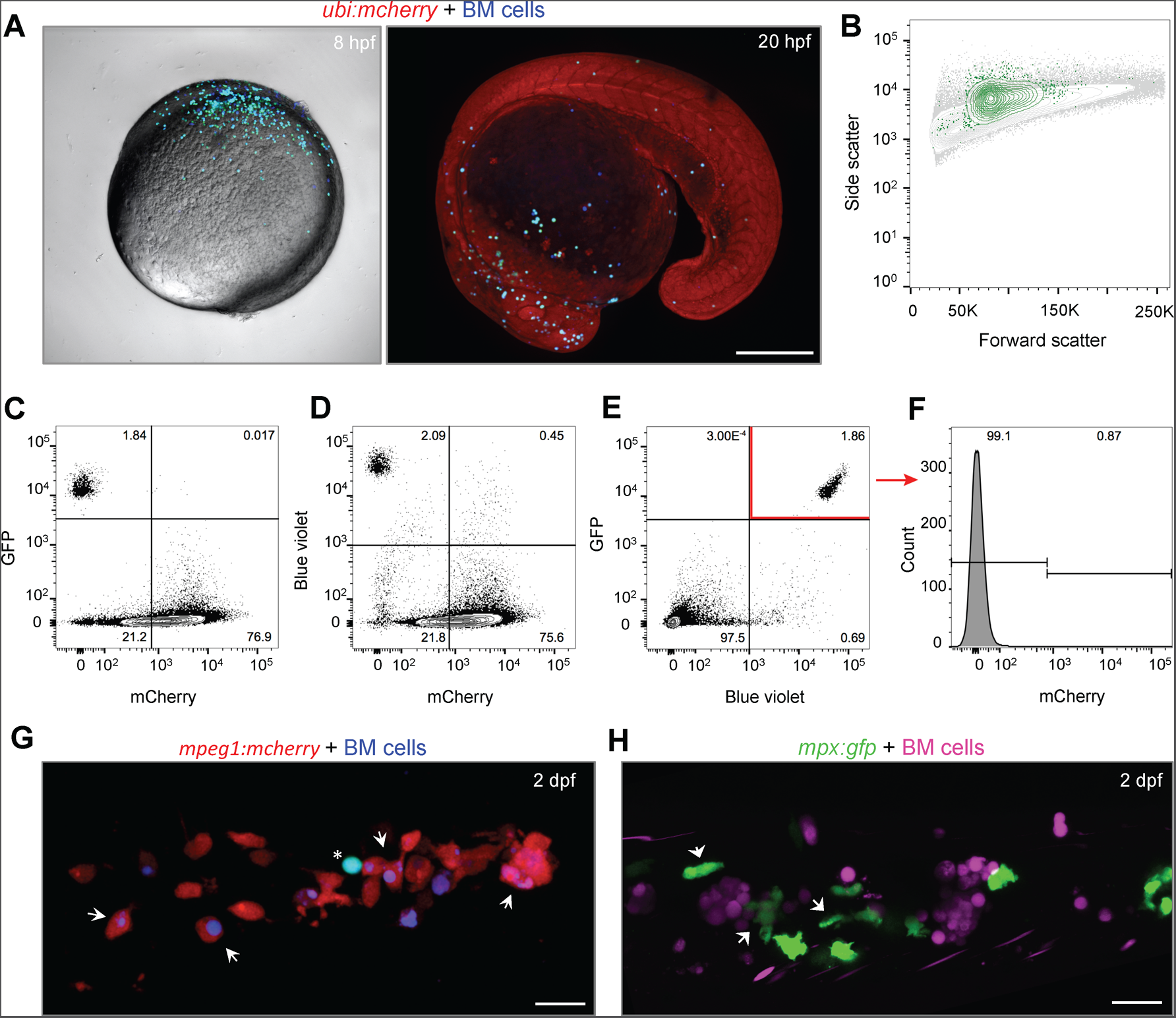
Xenotransplantation of mouse cells into zebrafish blastulae does not result in cell fusion events or vital dye transfer. A) Murine bone marrow cells (UBI-GFP) were blue labelled and transplanted into *ubi:mcherry* transgenic zebrafish blastulae. Confocal images from an 8 hpf (left panel) and a 20 hpf (right panel) transplanted embryos show no triple labeled cells (red, green and blue; white emission). Scale bar: 250 µm. B) At 2 dpf, 12 transplanted embryos were selected and a whole embryo cell suspension was prepared and analyzed by flow cytometry. Contour plot of the physical parameters identified by forward and side scatter show back-gated mouse bone marrow cells (green events) and fish cells (grey events). C) Contour plots from color based analysis show 0.017% of events as GFP+ and mCherry+, D) 0.45% of events as blue violet+ cells and mCherry+, and E) the majority of GFP+ as blue violet+. F) mCherry+ histogram plot of blue violet+/GFP+ pre-gated events (red arrow in D) showing 0.87% of triple labeled cells. G) Murine bone marrow cells (UBI-GFP) cells were blue labelled and transplanted into *ubi:mcherry* transgenic zebrafish blastulae. Confocal z-stack image of the fish PBI showing macrophage (red labeled) internalization of mouse blue+/GFP-cells (white arrows). Asterisk indicates double blue and green labeled cell (cyan). Scale bar: 20 µm. H) Confocal pseudocolored z-stack image of the fish PBI from a *mpx:gfp* fish transplanted with blue labelled murine bone marrow cells showing activated neutrophil cells (white arrows) around murine cells (magenta). BM cells: mouse bone marrow cells. Scale bar: 20 µm.

Nonetheless, we wished to evaluate the origin of double and triple labeled cells. To ascertain if endogenous cell-phagocytosis of murine cells could explain these events, we analyzed innate immune cell responses in the chimeras. First, we transplanted blue-stained bone marrow cells from the UBI-GFP transgenic mice into the macrophage labeled *mpeg1:mcherry*^33^ transgenic reporter fish. We found that endogenous hematopoiesis is not affected despite the presence of mammalian cells. However, live imaging showed that some fish macrophages residing in the CHT contained blue, but not green, labeled cell material (Figure 2G). This suggests that endogenous immune cells may recognize and phagocytose murine cell debris or dying murine cells that have lost GFP expression. We next transplanted blue stained bone marrow cells into the neutrophil labeled *mpx:gfp*^34^ transgenic reporter fish. As before, neutrophil development proceeded normally. A fraction of the fish neutrophils displayed an activated morphology in the chimeras, but showed no internalization of murine cells (~100 larvae) (Figure 2H). These results suggest macrophages, but not neutrophils, phagocytose dying murine engrafted cells.

We next evaluated the extent to which murine xenotransplanted cells underwent cell death. Analysis of cell death markers in the chimeras at 2 dpf showed that only 5.4% of xenografted cells were TUNEL positive (Supplemental Figure S12). This result suggests that there are few dying murine cells in the chimeras, which is in agreement with the flow cytometry analyses that detected less than 1% double or triple labeled cells. Furthermore, even if xenotransplanted cells were recognized as foreign by the zebrafish innate immune system, live murine cells can be tracked inside chimeric fish larvae for up to 6 dpf (Supplemental Figure S13). Overall, these results indicate that, with our procedure, a high number of murine cells can be transplanted into zebrafish blastulae that remain viable during the early larval stage.

### Real-time tracking of murine bone marrow cells in zebrafish reveals heterogeneous cell behaviors

Using live imaging, we next analyzed the behavior of the xenotransplanted murine bone marrow cells as zebrafish development proceeds. Visualization of chimeric animals at around 20 hpf showed murine cells distributed and moving over the yolk sac along the migratory route of endogenous primitive macrophages^39^ (Figure 3A and Supplemental Video SV1). Visualization of animals at 1 dpf showed that a subset of murine cells had entered the fish circulatory system, a phenomenon observed regardless of the total xenografted cell number. At 2 dpf, murine cells could be observed circulating within the entire fish bloodstream (Figure 3B and Supplemental Video SV2). Visualization at higher magnification revealed that circulating murine cells displayed variable velocities and adhesive rolling behaviors, indicative of differential adherence to the vascular endothelium (Figure 3C–C’ and Supplemental Videos SV3–SV4). Similarly, visualization of murine and fish cells within the CHT showed highly motile behaviors, which suggest dynamic niche interactions (Figure 3D–E and Supplemental Videos SV5 and SV6). Overall, these results suggest that xenografted murine cells have heterogeneous phenotypes and display different behaviors in the zebrafish chimeras.

**Figure 3.**
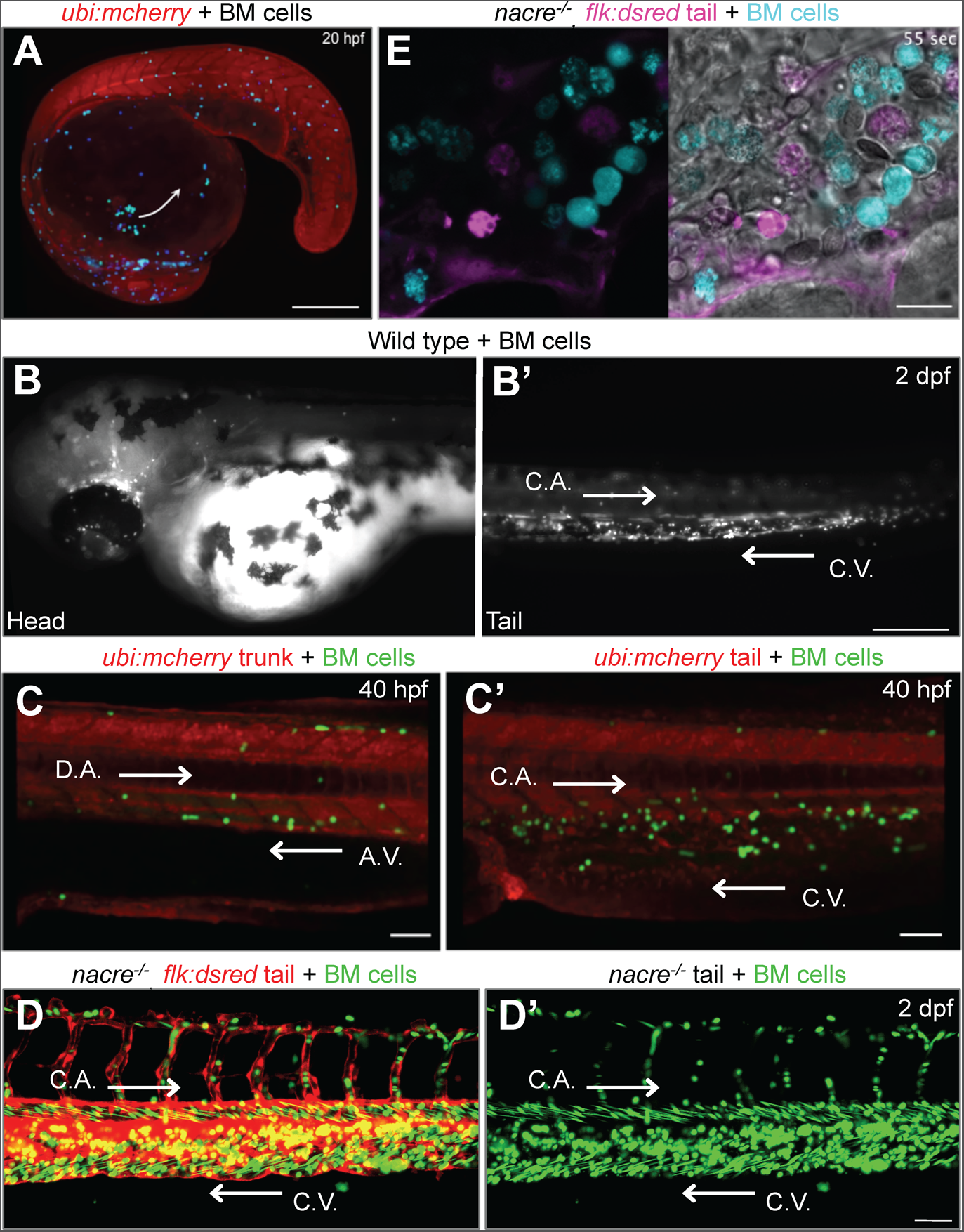
Live imaging of xenotransplanted zebrafish embryos and larvae shows migration and behavior of mouse bone marrow cells. A) Confocal image obtained from a 13 hour timelapse sequence that shows mouse bone marrow cells migrating along the route of endogenous primitive macrophages. Arrow depicts cell migration direction over the yolk sac (see Supplemental Video S1). Scale bar: 200 µm. B–B’) Epifluorescence images obtained from a time-lapse sequence that show mouse cells circulating within the B) fish head and B’) tail vasculature (see Supplemental Video S2). Scale bar: 250 µm. C–C’) Confocal images from a time-lapse sequence showing ubi-GFP transgenic mouse cells (green) interacting with fish endothelial cells within C) the fish trunk dorsal aorta and axial vein, and C’) the tail caudal aorta and caudal vein in an *ubi:mcherry* fish (see Supplemental Videos S3 and S4). Scale bar: 50 µm. D-D’) Pseudo-colored confocal images captured from a 1.5 hours time-lapse sequence showing green mouse cell dynamics within the fish caudal hematopoietic tissue in a vasculature reporter *flk:dsred* animal. Panel D’ is the green channel of panel D (see Supplemental Video S5). Scale bars: 50 µm. E) Pseudo-colored confocal image from a time-lapse sequence that shows individual mouse cell (cyan) dynamics at high magnification inside the fish CHT in a *flk:dsred* animal (see Supplemental Video S6). Scale bar: 10 µm. C.A.: caudal aorta; C.V.: caudal vein; D.A.: dorsal aorta; A.V.: axial vein. White arrows in B, C, C’, D and D’ depict fish blood flow direction.

### Xenotransplanted murine bone marrow cells respond to bacterial infection in fish

We had observed that murine bone marrow derived cells xenografted into zebrafish can express myeloid terminal differentiation markers. In order to determine if these cells are able to functionally respond to an inflammatory signal, 3 dpf chimeric larvae were infected with a clinical isolate of *Klebsiella pneumoniae,* an important human bacterial pathogen^40^. Intramuscular injection with ~100 cfu of *K. pneumoniae* elicited changes in the distribution of murine cells within the CHT compared to non-injected fish. Five hours after bacterial infection, murine cells became depleted from the hematopoietic niche (Figure 4A–B’). Intramuscular injection with a more concentrated inoculum of bacteria (~400 cfu) resulted in murine cells accumulating at the infection site starting at 1 hour post infection (hpi), with increased numbers at 24 hpi (Supplemental Figure S14). These experiments reveal that murine cells change their distribution in zebrafish chimeras upon bacterial infection and suggest they are mounting a specific response to this stimulus.

To visualize the interaction dynamics between murine cells and bacteria *in vivo*, we injected ~500 cfu of live red fluorescent *K. pneumoniae* into the otic vesicle of 2 dpf xenotransplanted *flk:dsred* larvae, and performed time-lapse imaging starting at 9 hpi. Live imaging showed murine cells migrating and interacting with bacteria, with some murine cells showing intracellular bacterial load (Figure 4C–G and Supplemental Video SV8). These results indicate that murine cells are able to detect and interact with bacterial cells in a zebrafish.

**Figure 4.**
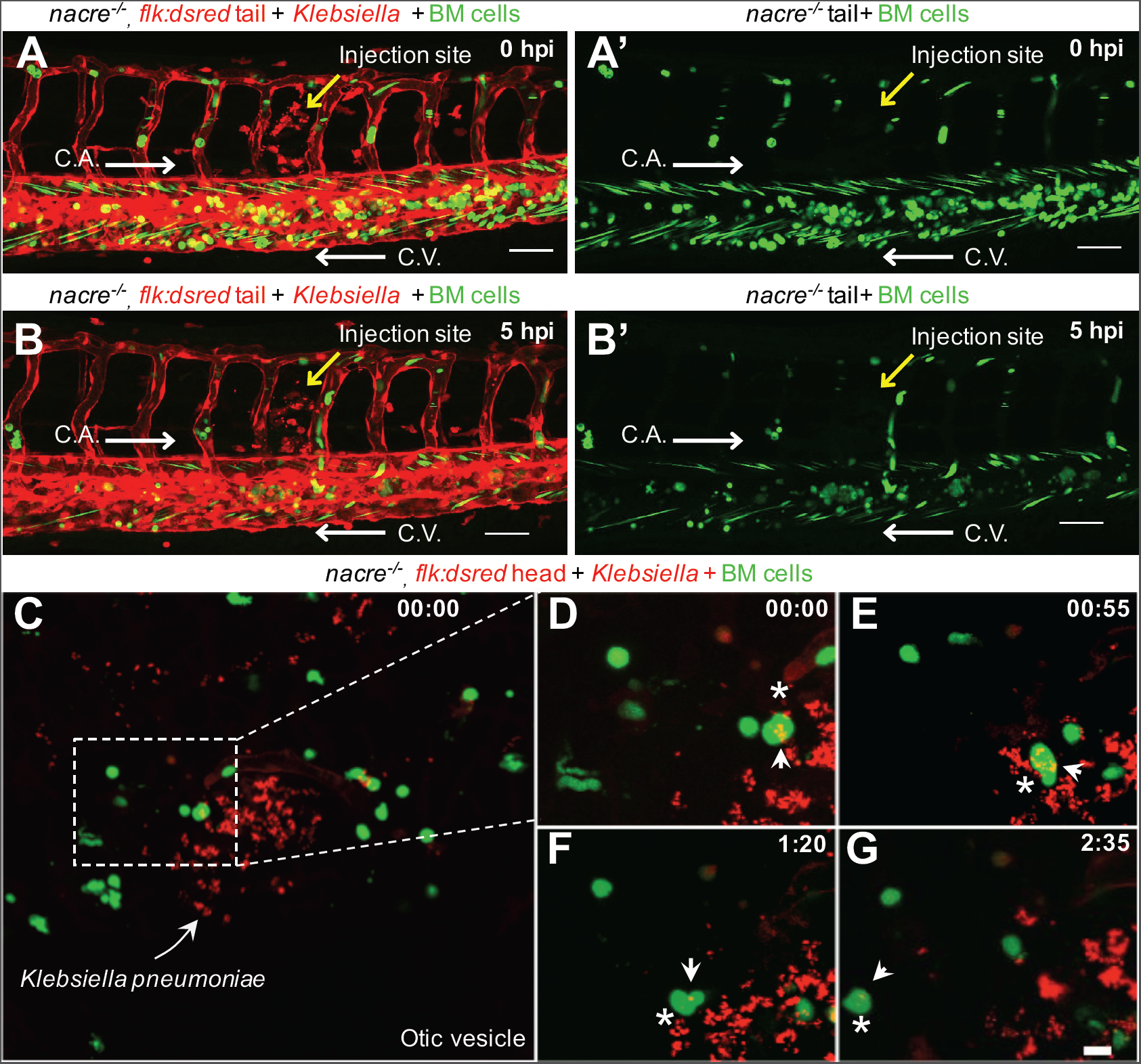
Xenotransplanted mouse cells respond to a bacterial infection in zebrafish. Pseudo-colored confocal images from a time-lapse sequence of xenotransplanted *flk:dsred* fish infected with *K. pneumoniae*. Endothelial cells are red-labeled, mouse cells are green-labeled and bacteria are red-labeled. A, A’–B, B’) Images acquired right after (A, A’; 0 hpi), or 5 hours after (B, B’; 5 hpi) intramuscular infection with ~100 cfu of *K. pneumoniae*. (A’–B’) Green channel reveals depletion of mouse cells in the CHT at 5 hpi (see Supplemental Video S7). Scale bars: 50 µm. (C–G) Confocal images captured from a time-lapse sequence centered on the otic vesicle of a fish infected with ~500 cfu of *K. pneumoniae*. Arrow points to red-labeled bacterial cells at the injection site. (D–G) Enlarged images showing the interaction of a single mouse cell (asterisk) with bacteria (arrow) that was tracked during the indicated times; the mouse cell containing bacteria later migrates away from the infection site (see Supplemental Video S8). Scale bar: 10 µm. C.A.: caudal aorta; C.V.: caudal vein; cfu: colony-forming units; hpi: hours post infection. White arrows in a, a’, b and b’ depict fish blood flow direction. Yellow arrow in a, a’, b and b’ depicts bacterial microinjection site.

## DISCUSSION

The transparency and tolerance to experimental manipulation of zebrafish embryos has extended their use in unanticipated ways, which include the transplantation of animal, plant and bacterial cells^8–27, 47–48^. For instance, a wide body of work has shown that xenotransplantation of human cancer cells into zebrafish is a useful *in vivo* platform to assess and screen for molecules controlling cancer cell metastasis and migration. Until now though, zebrafish xenotransplantation assays to visualize and study normal mammalian hematopoietic or immune cell development and function in living fish embryos have not been established. To broaden the scope of available cell xenotransplantation assays in zebrafish we have developed a method that generates zebrafish embryos with transiently chimeric hematopoietic tissue. This procedure constitutes a simple and efficient technique that allows the engraftment of mammalian hematopoietic cells into the fish embryonic developmental hematopoietic program. As a result, mammalian cells integrate into the developing primitive and definitive hematopoietic tissues of the host, and chimeric animals grow with a heterogeneous population of mammalian bone marrow-derived cells amidst their own hematopoietic niche. The xenografted animals allow for a broad spectrum of *in vivo* experiments to be carried out readily without any further manipulations and to continually visualize the cell behaviors that might ensue.

The method developed here to generate mammalian-zebrafish chimeras differs from previously existing ones in three main aspects. First, this method increases by 10-fold the number of cells that can be xenotransplanted in zebrafish. Xenotransplantation into the fish bloodstream has been previously reported for a range of 50–500 cells ^25–26^. In contrast, injection into the animal pole of a blastula shows that up to 5,000 mammalian cells can be transplanted into a fish. Strikingly, the animals develop normally considering the substantial number of exogenous cells. Of note, this might be possible only for hematopoietic cells, since transplantation of similar numbers of other cell types (or transformed cells) results in developmental abnormalities and high lethality, possibly due to unregulated expression of morphogens or adhesion molecules that impair fish development. Second, this method broadens the range of potential experiments that can be performed in chimeric animals. Xenotransplantation of cells within the fish bloodstream leads to their localization within the vasculature of the hematopoietic niche^25–26^. In contrast, xenotransplantation in the blastula stage allows cells to be integrated into the bloodstream as well as the developing hematopoietic tissues. This may allow analysis of hematopoietic cell-niche interactions. It also allows to examine the process of homing that progenitors undergo to migrate specifically to the hematopoietic tissue. In fact, we noted that xenotransplanted cells followed the same path taken by endogenous hematopoietic progenitors, suggesting they are following the same guidance cues at the appropriate developmental time. Thirdly, this method expands the developmental time frame of experiments that can be conducted. Transplantation into the bloodstream typically occurs in 2 dpf animals, restricting experiments to later developmental stages. Transplantation into blastula stage embryos allows experiments to be started as soon as 6 hours post fertilization. In sum, our methodology improves and broadens zebrafish blood cell xenotransplantation studies.

Monitoring of zebrafish chimeras throughout embryonic development evidenced murine cells migrating along and co-localizing, in succession, to the fish yolk sac, ICM, AGM, PBI, and later, the CHT and developing thymic lobes. These results strongly suggest that murine cells are actively sensing and responding to molecular cues originating from both the host primitive and definitive hematopoietic niches. The low frequency of chimeric animals displaying murine cells localized to the thymic lobes suggests that only specific bone marrow cell subsets or derivatives may be able to colonize the thymus. These cells may be poorly represented in the cell populations recovered using our selection method and their identity needs to be further determined. Since endogenous HSPCs start colonizing the developing thymus and kidney marrow at 3 dpf in order to give rise to the definitive wave of hematopoiesis in zebrafish^41–42^, it would be interesting to evaluate if xenotransplanted cells localizing to the thymus also correspond to murine hematopoietic stem cells. We also observed zebrafish chimeric animals that displayed murine cells within the pronephric tubules and kidney rudiment at 3 dpf. Previous reports have described a migration pathway along the pronephric tubules that initiates adult hematopoiesis in the developing kidney^43^. However, in the chimeras, we cannot exclude that the label we observe in the kidney could be associated with elimination of murine-cell debris. Importantly, transplantation of murine neutrophils and human promyelocytic cells, but not human neuroblastoma and human epithelial cells, resulted in colonization of the CHT, suggesting cell lineage homing specificity. Altogether, these results demonstrate that mammalian hematopoietic cells can recapitulate early zebrafish hematopoiesis, implying high conservation of homing molecules during hematopoiesis among vertebrates. In addition, we demonstrate that zebrafish hematopoietic chimeric embryos can be used to study *in vivo* and in real time mammalian developmental hematopoietic trafficking events. While we did not follow the fate of murine cells after the sixth day post transplantation, it will be interesting to determine whether their continued survival extends this period of time and past the emergence of the fish adaptive immune system.

A closer evaluation of cells homing to and residing within the fish hematopoietic tissue in chimeras at 2–3 dpf, evidenced murine cells expressing antigens related to stem and progenitor cells as well as active cell proliferation. The proportion of c-kit+ cells in the CHT (~35%) is slightly lower than the ~50% of c-kit+ cells observed in the HSPCs-enriched bone marrow cell population prior to transplantation. In addition, murine cells display antigens related to myeloid lineage cells after arriving at the CHT. Further, when observing live xenotransplanted larvae, we saw cells in circulation, some of which displayed “rolling” behaviors, akin to what has been described for leukocyte-endothelium interactions. We hypothesize that proliferating murine hematopoietic progenitors differentiate into myeloid lineage cells in zebrafish, as has previously been reported for human CD34+ cells^25^. To what extent murine cell differentiation takes place in a zebrafish, and if this is a cell-autonomous process or host driven, needs to be further addressed. What other cell types are present or become engrafted in the chimeras needs to be further explored as well. Regardless, our results reveal the existence of different cell types in the chimeras, supporting the concept that zebrafish hematopoietic tissue chimeric animals can be used to study the cell biology of mammalian HSPCs and derived cells *in vivo*, as well as their interactions within the hematopoietic niche and endothelial vasculature.

The zebrafish is an excellent model for the study of innate immunity and of the inflammatory response elicited by tissue damage or infection^44–46^. Given that we had shown that murine HSPCs differentiate into neutrophils and macrophages in the fish, we wished to know whether they had acquired functional competence as well. Chimeric larvae challenged by localized intramuscular infection with a low dose of with *K. pneuomoniae* revealed a dramatic decrease of murine cell numbers in the fish CHT, with an increase in murine cell numbers in the vessel directly proximal to the inoculum injection site. This suggests that murine cells respond to the infectious challenge by entering the circulation and homing to the bacterial site. However, we cannot exclude the possibility that part of the observed effect is due to activation of endogenous immune cells or emergency granulopoiesis that might lead to elimination of the murine cells in the CHT. Infection with a higher inoculum of bacteria (~400 cfu) resulted in specific localization of murine cells near the injection site starting as early as 1 hpi, with increased numbers of infiltrated cells over time. These results suggest murine cells actively migrate towards the infection site, although bacterial-induced tissue necrosis might also favor the extravasation or interstitial migration of murine cells towards the infected site. In addition, murine cells can be observed actively interacting with bacterial cells when these have been injected into the otic vesicle, with some murine cells evidencing an intracellular bacterial load indicative of phagocytosis. It remains to be determined what murine immune cell types are manifesting these responses, as well to characterize which molecules they are expressing when stimulated by bacterial cells in a zebrafish. Nevertheless, our results show that xenografted murine cells are able to respond to an infection and we demonstrate, for the first time, that zebrafish chimeric animals can be utilized to study mammalian immune cell host-pathogen interactions *in vivo*.

In conclusion, our results introduce the use of murine-zebrafish hematopoietic tissue chimeras for studying murine bone marrow derived cell dynamics and their interaction within stromal niches and pathogens *in vivo*. Both murine and human hematopoietic derived cells can be engrafted into the fish CHT. Therefore, the range of potentially valuable applications of this simple yet powerful technique can include diverse cell types and species. Future improvements could incorporate the use of immunodeficient zebrafish hosts to extend the survival of the xenografted cells as has been shown in humanized mice^1^; this could also benefit the analysis of mammalian cell responses without the interference of competing endogenous cells. In addition, the utilization of murine transgenic reporter lines (*e.g.* UBI-GFP) could allow tracking of murine cells for more extended periods of time. We envision that zebrafish hematopoietic tissue chimeric embryos could be adapted to study *in vivo* and in real time the contribution of cell-autonomous versus non-cell autonomous gene functions in diverse processes such as homing mechanisms, tolerization and host-pathogen interactions, among others. Ultimately, the efficiency of transplantation achieved with this technique could allow targeted drug screens or even patient-specific assays to be carried out rapidly in an *in vivo* experimental setting^49–50^.

## ACKNOWLEDGMENTS

Funding for this work was provided by FONDAP (15090007), FONDECYT (1140702) to M.L.A., and MECESUP (UCH 0713) to M.P. We thank Travis Walton for his helpful comments on the manuscript.

## AUTHORSHIP

M.P., E.J.V. and M.L.A. conceived the idea. M.P., A.C., C.P., C.E., H.J, E.J.V., D.H. and M.L.A. designed or performed transplantation and infection experiments. M.P, E.J.H., C.P., A.N., J.E.H., B.L, R.R and L.I.Z. designed or conducted murine cell behavior analyses. M.P. and M.L.A. wrote the manuscript. M.P., A.C., E.J.H, C.P., B.L., E.J.V., L.I.Z, D.H. and M.L.A. performed data analysis and interpretation.

